# Linezolid Acts as a Selective Inhibitor of the JAK2^V617F^ Mutation

**DOI:** 10.64898/2026.05.31.729063

**Authors:** Anisa Azatovna Gumerova, Christoph Schaniel, Zixing Huang, Althea Agdamag, Shuhui Liu, Alessandro Principi, Jacob Kazmi, Francisco Fueyo Gonzalez, Jihong Cui, Georgii Pevnev, Caliana Yang, Zehra Tumoglu, Xin Gao, Tony Yuen, Yelena Ginzburg, Jeffrey Glassberg, Shozeb Haider, Mone Zaidi, Ronald Hoffman, Huihui Li

## Abstract

The *JAK2*^V617F^ (JAK2^VF^) driver mutation is found in 95% of patients with polycythemia vera (PV), a progressive myeloproliferative neoplasm. Current treatments suppress excessive hematopoiesis but lack specificity for targeting JAK2^VF^ cells, are unable to deplete mutant stem/progenitor cells and ultimately result in drug resistance. We discovered that the FDA-approved antibiotic, linezolid (LZD), ameliorates the PV phenotype across multiple model systems. LZD suppressed cell proliferation and STAT5 signaling, altered the cell cycle, and increased apoptosis of *JAK2*^VF^-harboring human erythroleukemia cells, but not in wild-type acute leukemia cells. Computational modelling indicated that LZD interacts specifically with mutant JAK2^VF^ but not with wild-type JAK2 protein. We further showed that, in *JAK2*^VF^ mice that faithfully recapitulate human PV, LZD mitigates disease burden by selectively targeting *JAK2*^VF^ stem cells thereby normalizing spleen size and blood counts. LZD also inhibited hematopoietic colony formation by patient-derived peripheral blood mononuclear cells, with the more primitive progenitors being preferred targets. Importantly, LZD selectively decreased JAK2^VF+^ colony numbers, without impacting wild-type JAK2 colonies. In all, the data provide a firm foundation for evaluating LZD-like molecules as an effective therapy for PV and other myeloproliferative neoplasms.

**Key points:**

1. Linezolid acts as a JAK2^V617F^IZselective inhibitor in PV mouse models and PV patient samples while sparing wildIZtype hematopoiesis.
2. Linezolid acts directly on JAK2^V617F^ hematopoietic stem cells.

## INTRODUCTION

Philadelphia chromosome-negative myeloproliferative neoplasms (MPNs) are a group of blood disorders that include polycythemia vera (PV), essential thrombocythemia (ET) and myelofibrosis (MF). MPNs are hematopoietic stem cell (HSC) disorders and are associated with chronic inflammation, abnormal myelopoiesis, and driver mutations in *JAK2*, *CALR* and *MPL*^1^. *JAK2^V617F^* (JAK2^VF^) is the most common driver mutation found in MPN patients with a frequency of ∼95% in PV and 50-60% in ET and MF. PV has a chronic, indolent clinical course and is characterized by an increase in red blood cell production as measured by elevated hemoglobin (Hb) levels and an increased risk of thrombosis^2^. However, approximately 13% of PV patients progress to a syndrome indistinguishable from overt forms of PMF, characterized by ineffective hematopoiesis causing cytopenias, increased marrow micro vessel density, and progressive marrow fibrosis resulting in extramedullary hematopoiesis and splenomegaly, with about 7% of patients eventually developing a secondary form of acute myeloid leukemia.

Currently available drug therapy is not curative. Allogeneic hematopoietic stem cell transplantation currently is the only curative option for MPN patients, albeit associated with high risk of morbidity and mortality. Current treatments for PV include phlebotomy to control hematocrit levels and cytoreductive therapy with hydroxyurea (HU), several interferon formulations and JAK2 inhibitors that aim to suppress hyperactive hematopoiesis^2–4^. However, these therapies often result in suppression of both malignant and healthy hematopoietic cells^3^ and intolerance or resistance to the therapeutic agent^4^, highlighting a significant unmet need for safer, more targeted therapies. Studies demonstrated that the MPN-propagating cells responsible for disease burdens are exclusively Jak2^VF^ HSC^4,5^.

A feature of MPNs is increased inflammatory cytokine elaboration, which correlates with worsening systemic symptom burden and disease prognosis^1^. The role of the microbiome in modulating inflammation in the context of cancer, including MPNs, is becoming increasingly obvious^6,7^. Linezolid (LZD), an FDA-approved antibiotic medication used to safely treat patients with serious gram-positive bacterial infections^8^.^8,9^. A few case reports demonstrated that some patients on LZD developed RBC aplasia/anemia and myelosuppression^10–15^, providing a rationale for investigating LZD as a suppressor of malignant hematopoiesis.

Here, we report our findings that LZD is a potent JAK2^VF^-selective inhibitor that is highly efficacious in reducing disease burden and targeting malignant HSCs, while sparing normal hematopoietic cells.

## METHODS

### Mice

Male and female C57BL/6J (CD45.2; RRID:IMSR_JAX:000664), B6.SJL-*Ptprc^a^Pepc^b^*/BoyJ (CD45.1, RRID:IMSR_JAX:002014), and C57BL/6-Tg(UBC-GFP)30Scha/J (*UBC-GFP*, RRID:IMSR_JAX:004353) mice aged 8 weeks were obtained from the Jackson Laboratory and housed under specific pathogen-free (SPF) conditions. Knock-in human *JAK2*^VF^ mice^16^ were obtained from the Albert Einstein College of Medicine, quarantined for 6 weeks before breeding in-house. Each line was bred in trio and the genotype of the offspring was confirmed by PCR on mouse tail genomic DNA. All procedures were performed in accordance with the Institutional Animal Care and Use Committee (IACUC) guidelines of Icahn School of Medicine at Mount Sinai. All experiments were carried out with 5 mice per group, and repeated 2–3 times, except for the experiment described in supplemental Figure 1, in which 3 mice per group were used.

### *In vivo* LZD treatments

Marrow-transplanted mice received drinking water containing 1 g/L linezolid (Chem-Impex, Cat# 29723-25G), which was replenished every other week for the duration of time mentioned in each experiment. Control/untreated group received regular water. At week 12 after bone marrow transplantation, the recipient mice were euthanized and body and organ weights measured. Peripheral blood, serum, bone marrow, and spleen cells were harvested and directly analyzed or cryopreserved either in liquid nitrogen or at −80°C.

### Hematopoietic Colony Assays

A total of 300 CD34+ cells were mixed with 1.1 mL of methylcellulose medium and plated in triplicate onto 35-mm culture dishes (StemCell Technologies). Cells were treated with 5, 10 or 50 µg/mL LZD. Three untreated plates were used as control. The plates were incubated at 37°C in a humidified 5% CO₂ incubator for 14 days. Cytokine-s upplemented cultures use IMDM-based methylcellulose medium containing erythropoietin (EPO, 1 U/mL), stem cell factor (SCF, 100 ng/mL), granulocyte-macrophage colony-stimulating factor (GM-CSF, 50 ng/mL), FMS-like tyrosine kinase 3 ligand (FLT3L, 50 ng/mL), thrombopoietin (TPO, 50 ng/mL), interleukin-6 (IL6, 50 ng/mL) and interleukin-3 (IL3, 25 ng/mL) to promote optimal colony growth.

Colonies were enumerated using an inverted phase-contrast microscope (Model CKX41, Olympus). Colonies were identified and quantified based on standard haematopoietic colony morphology as follows. Burst-Forming Unit-Erythroid (BFU-E) are large, multi-clustered colonies with a dispersed appearance. Colony-Forming Unit-Erythroid (CFU-E) are smaller, single-clustered colonies. Granulocyte-Macrophage Colony-Forming Units (CFU-GM) are compact colonies composed of granulocytes and macrophages. Granulocyte, erythrocyte, macrophage, megakaryocyte colony-forming unit- (CFU-GEMM) are mixed colonies containing a variety of blood cell types that can give rise to granulocytes, erythrocytes, macrophages and megakaryocytes. The number of each colony type were enumerated per culture plate.

### Hematopoietic cell lines utilized

Human erythroleukemia (HEL 92.1.7, Cat# TIB-180) and B-cell acute lymphoblastic leukemia (RS4;11, Cat# CRL-1873) cell lines were obtained from ATCC and cultured in RPMI-1640 medium (Gibco) supplemented with 10% foetal bovine serum (FBS; Gibco) and 1% penicillin–streptomycin (Gibco) at 37°C in a humidified incubator containing 5% CO₂. Cells were passaged when cultures reached approximately 70% confluency. HEL and RS4;11 cells were seeded at 1.75 x 10^5^ cells/well in separate 12-well plates, and were treated, in triplicates, with linezolid at concentrations of 0, 10, or 50 μg/mL. Cells were passaged on day 4. Cell number and viability were assessed on days 4 and 6 using acridine orange/propidium iodide (AO/PI) staining (Nexcelom Bioscience, Cat# CS20106) and quantified using a Cellometer Auto 2000 automated cell counter (Nexcelom).

### Computational *in silico* analysis

The wild-type (WT) JH2 subdomain model was built starting from the Mg-ATP bound crystal structure (PDB: 4FVQ)^17^. Three mutations introduced in the experimental structure for stability during crystallization were reverted (W659A back to W659, W777A back to W777, and F794H back to F794), using the PyMOL Mutagenesis Wizard^18^.

The V617F JH2 subdomain model was also built starting from the Mg-ATP bound crystal structure (PDB: 4FVR)^17^. In this case, only 2 mutations needed to be reverted (W777A reverted to W777 and F794H reverted to F794).

Ligand docking was carried out on the JH2 subdomain (WT and V617F) monomers. Linezolid (LZD) was docked to the V617F monomer using ICM-Pro^19^. For the V617F monomer, the binding site was chosen using the ICM Pocket Finder tool with a low tolerance value (tolerance 1). A pocket (Volume 413.9 Å) was identified around the phenylalanine triad (F594, F595, F617) and used to dock the three compounds. Each ligand was docked following the ICM-Pro default docking procedure. The best pose was chosen based on the lowest ICM score. Since the binding pocket does not exist in the WT, the ICM Pocket Finder tool could not find the proposed binding site. In WT JH2 monomer, LZD was not expected to bind to the WT JAK2.

### Human Samples

De-identified male and female Jak2^V617F^ patient peripheral blood mononuclear cell samples were obtained from the Hematological Malignancy Tissue Bank at the Icahn School of Medicine at Mount Sinai, which stores and disburses specimens under patient-signed informed written consent approved by the Institutional Review Board at Mount Sinai. Six individual patient samples were assayed (supplemental Table 1).

Details on bone marrow transplantation, flow cytometry, hematological analyses, histology, RNA isolation and quantitative real-time PCR, animal randomization and statistics, and data collection and availability are described in Supplemental data.

## RESULT

### LZD Rescues the PV Phenotype in *JAK2*^VF^ Mice and Has No Impact on WT Mice

The *JAK2*^VF^ mouse model of PV is characterized by erythrocytosis, leukocytosis, splenomegaly, and iron deficiency^16^. A cohort of such mice was generated by transplanting homozygous *JAK2*^VF^ mouse bone marrow into lethally irradiated WT recipients. While investigating the effect of the microbiome in this *JAK2*^VF^ mouse model, we unexpectedly found that mice treated with LZD (1 g/L in drinking water for 6 weeks) normalized their peripheral blood parameters with decreased red blood cell counts, hematocrit levels, reticulocytes, and white blood cell counts (Figure 1A) with no effect on mean corpuscular vome, mean corpuscular hemoglobin, or red cell distribution width (Figure 1B). These data suggest that LZD can rescue the PV phenotype in *JAK2*^VF^ mice, without changes in the appearance of JAK2^VF^ red blood cells. It should be noted that, as a negative control, the absence of a morphological change shows that LZD does not convert *JAK2*^VF^ red blood cells fully into WT cells, and therefore, as expected, does not affect variant allele frequency in the *JAK2*^VF^ mouse. Within 6 weeks of treatment, hematocrit levels dropped from 77% to 51% (Figure 1C) and splenomegaly was eliminated (Figure 1D), but bone marrow hypercellularity remained unchanged (Figure 1E). Interestingly, LZD-treated *JAK2*^VF^ mice displayed an increase in body weight as evidence for a reduced disease burden (Figure 1F).

**Figure 1:**
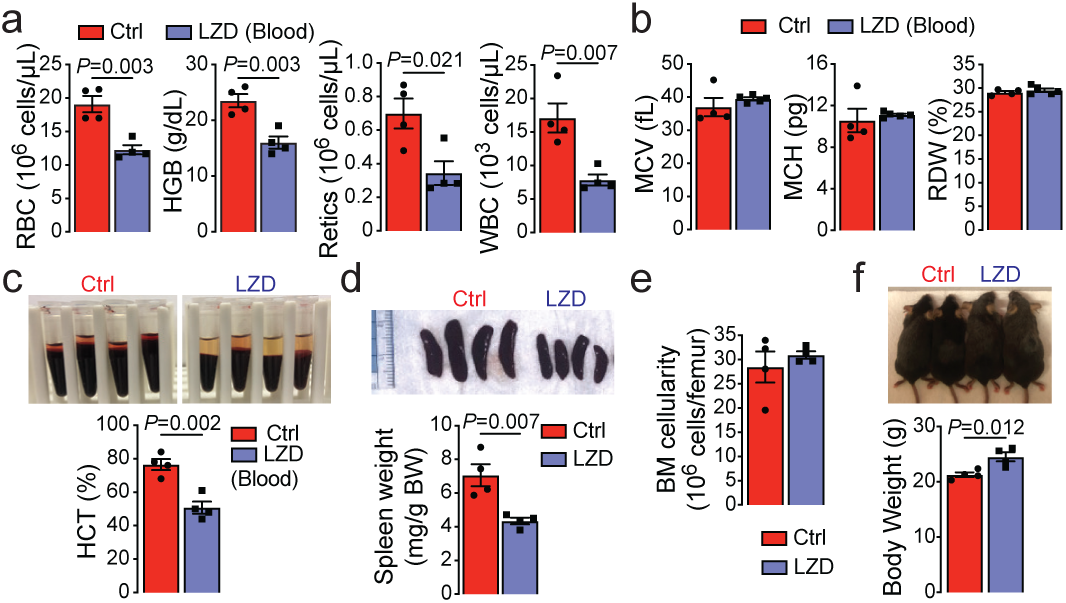
LZD normalized PV-like abnormalities in *Jak2*^VF^ mice. 3-month-old male homozygous *JAK2*^VF^ mice were treated with LZD (1 g/L in drinking water) for 6 weeks. Shown are peripheral blood cell counts, namely red blood cell (RBC), haemoglobin (HGB), reticulocytes (Retics) and white blood cell (WBC) (**a**); mean corpuscular volume (MCV), mean corpuscular hemoglobin (MCH) and red cell distribution width (RDW) (**b**); haematocrit (HCT) (**c**); spleen weight (**d**); bone marrow cellularity (**e**); and body weight (**f**) of *JAK2*^VF^ mice after 6 weeks of treatment. Mice receiving drinking water without LZD served as control (Ctrl). Statistics: mean ± s.e.m., *N*=4 mice per group, unpaired two-tailed Student’s *t*-test, *P* values shown.

We demonstrated that LZD treatment reduced the total Ter119+ erythroid cell numbers in both the bone marrow and spleen (Figure 2A). Furthermore, the percentage of enucleated red blood cells (Ter119^+^, FSC^low^, CD44^medium/low^) was decreased in the bone marrow with LZD treatment, and both erythroblasts (Ter119+, FSC^hi^, CD44^hi^). Furthermore, enucleated red blood cells were reduced in LZD-treated spleens (Figure 2B), which accompanied a reduced spleen size. In addition, LZD promoted the death of erythroblasts (DAPI+ cells) (Figure 2C). Elevated erythropoiesis in PV requires the increased availability of iron for hemoglobin synthesis resulting in the depletion of iron stores in the bone marrow and spleen^16,20^. LZD treatment normalized iron stores and suppressed CD71+ erythroblast numbers in bone marrow and spleen (Figure 2D, E). Morphological analysis further showed greater red pulp area compared to white pulp area in the spleens of untreated *JAK2*^VF^ mice, which was again normalized by LZD (Figure 2D). These findings provide clear evidence that LZD suppresses the accelerated hematopoiesis, reduces the splenomegaly, and normalizes iron utilization in PV mice.

**Figure 2:**
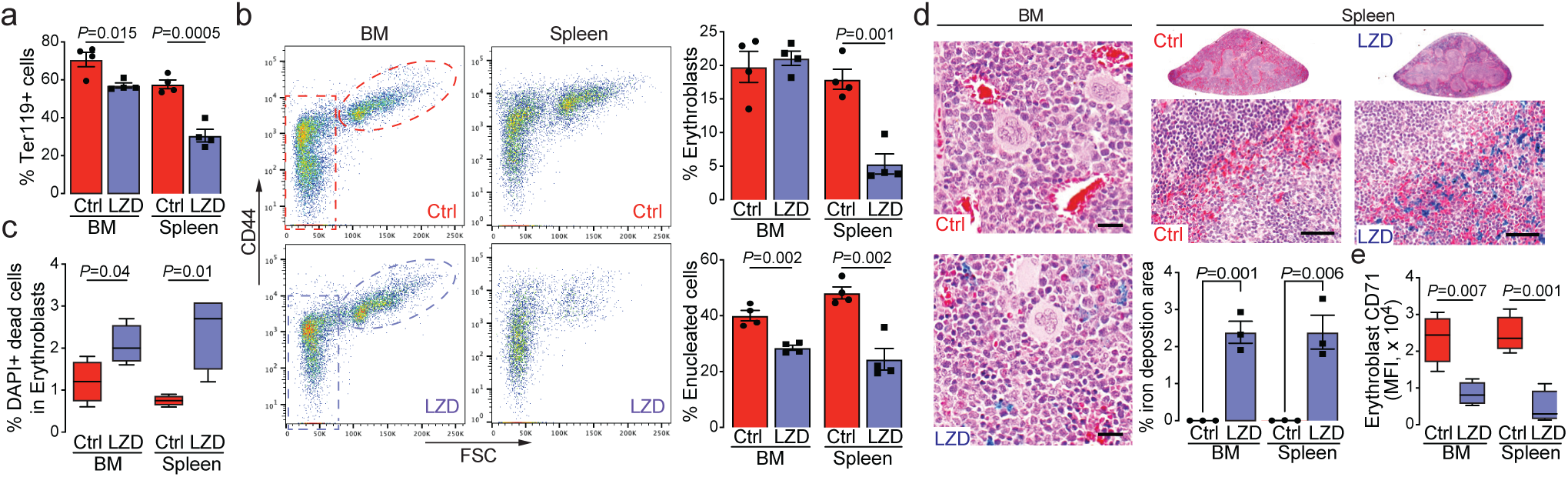
LZD reduced erythropoiesis and restored iron deposition in *JAK2*^VF^ mice. 3-month-old male homozygous *JAK2*^VF^ mice were treated with LZD (1 g/L in drinking water) for 6 weeks. Shown are total erythroid (% Ter119+) (**a**); erythroblasts (CD44^hi^ FSC^hi^, oval circle) and enucleated cells (CD44^medium^ FSC^low^, rectangle) (**b**); dead erythroblasts (DAPI+) (**c**); red/white pulp ratio (H&E) and % iron deposition area (Prussian blue) (**d**); and erythroblast transferrin receptor 1 (CD71) expression (mean fluorescence intensity, MFI) (**e**) in bone marrow (BM) and spleen of control and LZD-treated *JAK2*^VF^ mice. Scale bar (d): 20 µm (bone marrow), 50 µm (spleen). Statistics: mean ± s.e.m., *N*=4 mice per group, unpaired two-tailed Student’s *t*-test, *P* values shown.

By contrast, WT mice treated with the same dose of LZD as *JAK2*^VF^ mice showed a paradoxical, small but significant increase in red blood cell counts and hemoglobin levels, without changes in other hematological parameters (Figure 3A-C). The total percentage of erythroid cells was upregulated in the bone marrow of LZD-treated WT mice (Figure 3D, E). Similar to observations in *JAK2*^VF^ mice, LZD decreased CD71+ cell levels in the bone marrow of WT mice (Figure 3F). The mild increase in erythropoiesis in LZD treated WT mice may be due to a mild suppression of CD71 expression, particularly given that CD71 haploinsufficiency in mice is known to increase erythropoiesis^21,22^. Overall, however, the data collectively document that, unlike the potent therapeutic inhibition of hematopoiesis seen in *JAK2*^VF^ mice, LZD treatment of WT mice has a minimal impact.

**Figure 3:**
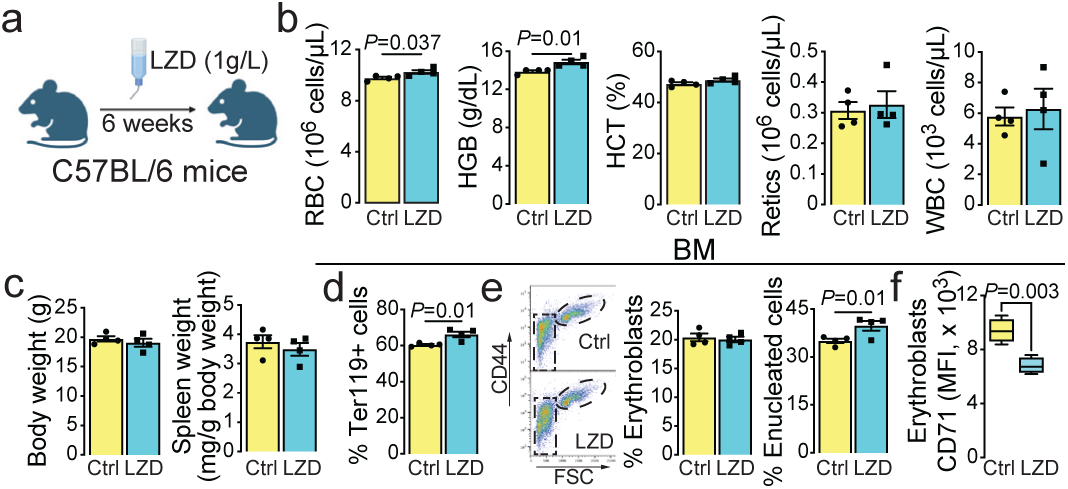
LZD treatment had minimal effects on *Jak2* wild-type mice. 3-month-old male C57BL/6 mice were treated with LZD (1 g/L in drinking water) for 6 weeks (**a**). Shown are peripheral blood counts, namely red blood cell (RBC), hemoglobin (HGB), hematocrit (HCT), reticulocytes (Retics) and white blood cell (WBC) (**b**) and body weight and spleen weight (**c**) of control (Ctrl) and LZD-treated C57BL/6 mice. Also shown are total erythroid cells (Ter119+) (**d**), erythroblasts (CD44^hi^ FSC^hi^, oval circle) and enucleated cells (CD44^medium^ FSC^low^, rectangle) (**e**) and erythroblast transferrin receptor 1 (CD71) expression (**f**) in bone marrow (BM). Statistics: mean ± s.e.m., *N*=4 mice per group, unpaired two-tailed Student’s *t*-test, *P* values shown.

### LZD Selectively Binds to Mutant JAK2^VF^ but Not to WT JAK2

To gain insight into the mechanism by which LZD causes amelioration of the PV phenotypes, we explored whether LZD differentially influences signaling pathways downstream of JAK2^VF^ mutation using a human erythroleukemia cell line, HEL, displaying homozygosity of JAK2^VF^, and the human acute leukemia cell line, RS4;11, with WT JAK2^23^. LZD significantly suppressed p-STAT5 signaling in HEL cells without affecting p-STAT3 and p-ERK (Figure 4A). This showed that LZD acts primarily by disrupting JAK2^VF^/STAT5 signaling. Cell cycle analysis of LZD-treated HEL cells further revealed fewer cells in G0, an accumulation of cells in G1, and a slight decrease in cells in S/G2/M at higher drug concentrations (Figure 4B)—indicating that LZD promotes the exit of cells from their quiescent state. LZD also increased apoptosis of HEL cells in a concentration-dependent manner (Figure 4C). As a result, LZD progressively inhibited the growth of HEL cells in a concentration-dependent manner between days 4 and 6 of incubation (Figure 4D). In clear contrast, LZD treatment of RS4;11 cells did not alter JAK-STAT signaling (Figure 4E). While LZD induced an increase in cells in G0 at the highest dose, it did not change the number of cells in G1 or S/G2/M phase, those undergoing apoptosis, or total cell numbers (Figure 4F-H). As a negative control, we evaluated the effect of ruxolitinib, a JAK1/2 inhibitor on HEL and RS4;11 cell lines. Ruxolitinib, which is used widely to treat PV patients, suppressed cell proliferation and promoted apoptosis in both HEL and RS4;11 cell cultures in a concentration-dependent manner (supplemental Figure 1A, B). These data provide compelling evidence that LZD selectively inhibits cells harboring the JAK2^VF^ mutation by affecting STAT5 signaling and altering cell cycling, while having limited to no effects on WT cells lacking this mutation.

**Figure 4:**
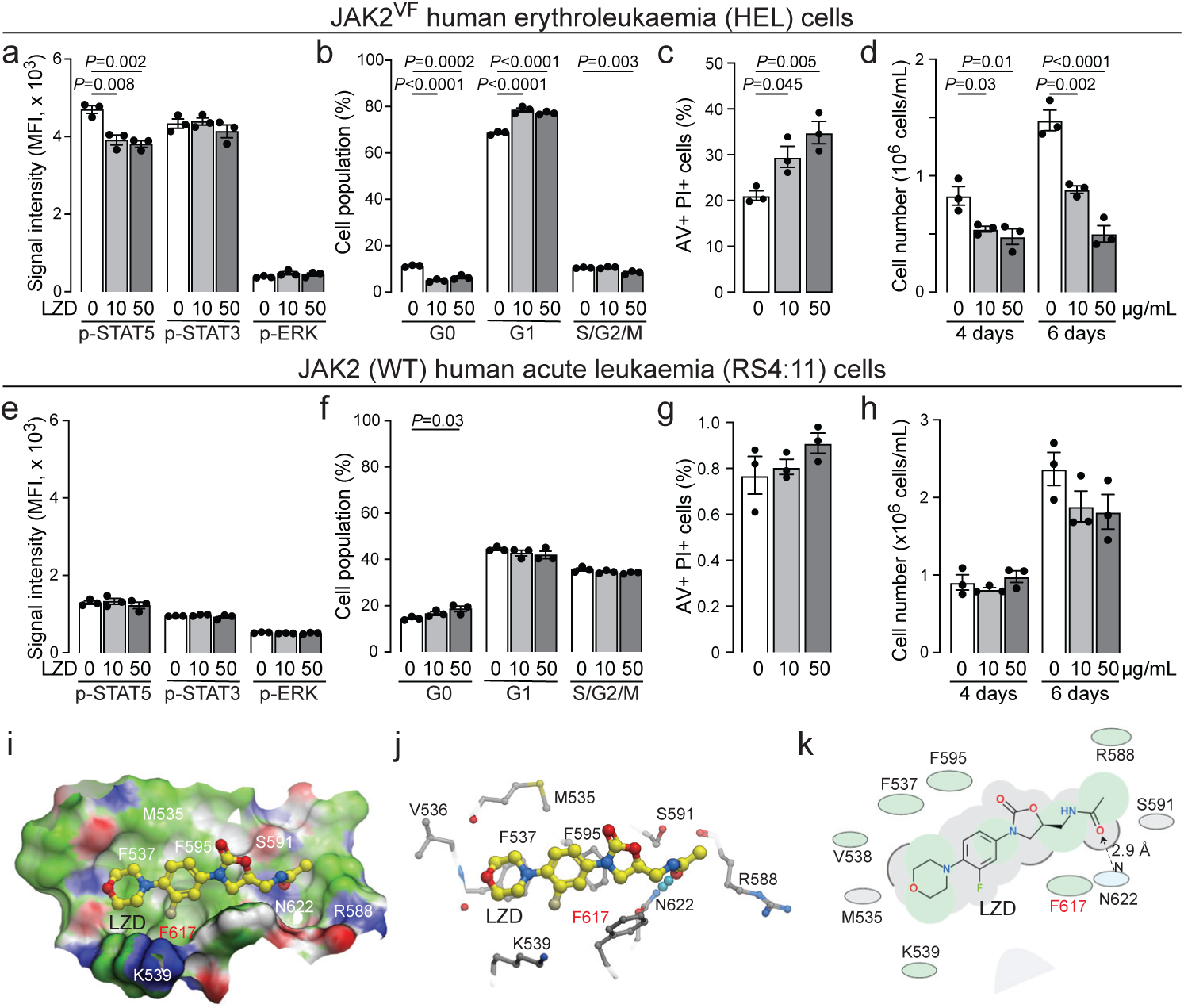
LZD selectively binds to JAK2^VF^ to reduce STAT5 phosphorylation, alter cell cycle distribution, induce apoptosis and inhibit cell growth. Shown are JAK2-STAT signaling (**a, e**), cell cycle (**b, f**), apoptosis (annexin V+ and propidium iodide+, AV+ PI+) (**c, g**) and cell growth (**d, h**) following 0, 10 or 50 µg/mL LZD treatment in JAK2^VF^ mutant HEL and JAK2 wild-type RS4:11 cells, respectively. Molecular docking of LZD onto JAK2^VF^ JH2 domain (space-filling model, **i**; ball-and-stick model, **j**; and 2D ligand interaction map, **k**) reveals a hydrogen bond between the acetamide oxygen of LZD and JAK2^VF^ N622 (dashed arrow, k), suggesting a strong, stabilizing interaction. Hydrophobic residues (V536, F537, F595, F617, and M535) surrounding the morpholine and phenyl rings of LZD form a complementary pocket that stabilizes the drug through non-polar interactions. Polar and charged residues (R588, K539 and S591) further stabilize ligand binding. Statistics (a–h): mean ± s.e.m., *N*=3 per group, unpaired two-tailed Student’s *t*-test (vs. 0 µg/mL LZD), *P* values shown.

To provide granular detail on the interaction of LZD with the JAK2VF mutation, we modeled JAK2VF and WT JAK2 crystal structures and docked LZD into their JAK2 pseudokinase domains (JH2), where the V617F mutation is located. The JH2 domain of JAK2^VF^ possesses a shallow ligand-binding pocket characterized by a complex electrostatic environment. The LZD-JAK2^VF^ JH2 complex exhibited a distinct energetic profile. Namely, the aromatic residues F617 (–1.74 kcal.mol⁻¹), F537 (–1.11 kcal.mol⁻¹), and F595 (–0.87 kcal.mol⁻¹) formed a π-stacking “sandwich”, significantly reducing adjacent polar penalties. Furthermore, residue N622 shifted toward a favorable interaction (–1.51 kcal.mol⁻¹) due to hydrogen bond formation with LZD (Figure 5I-K, supplemental Figure 3C). Together, these favorable interactions contributed to an overall binding free energy of –13.11 kcal.mol⁻¹ for LZD-JAK2^VF^ JH2 (supplemental Figure 1C). As a negative control, we analyzed the interaction of the JH2 domain with ruxolitinib, a JAK2 inhibitor that is known to have limited interactions with the JH2 domain^24^. We found consistently that substantial unfavorable contributions in the ruxolitinib-JAK2^VF^ JH2 complex originated from residues S591 (+4.72 kcal.mol⁻¹) and N622 (+2.43 kcal.mol⁻¹) (supplemental Figure 1D). Favorable compensation was limited and mainly provided by residue F617 (–2.28 kcal.mol⁻¹), ultimately leading to an overall unfavorable binding free energy (+2.05 kcal.mol⁻¹) (supplemental Figure 1D). Importantly, a docking site was not detected between LZD and WT JAK2 JH2 protein (supplemental Figure 1E). In all, the *in vitro* and *in silico* studies together provide foundational evidence for a highly selective interaction between the antibiotic LZD and the driver mutation in PV patients.

**Figure 5:**
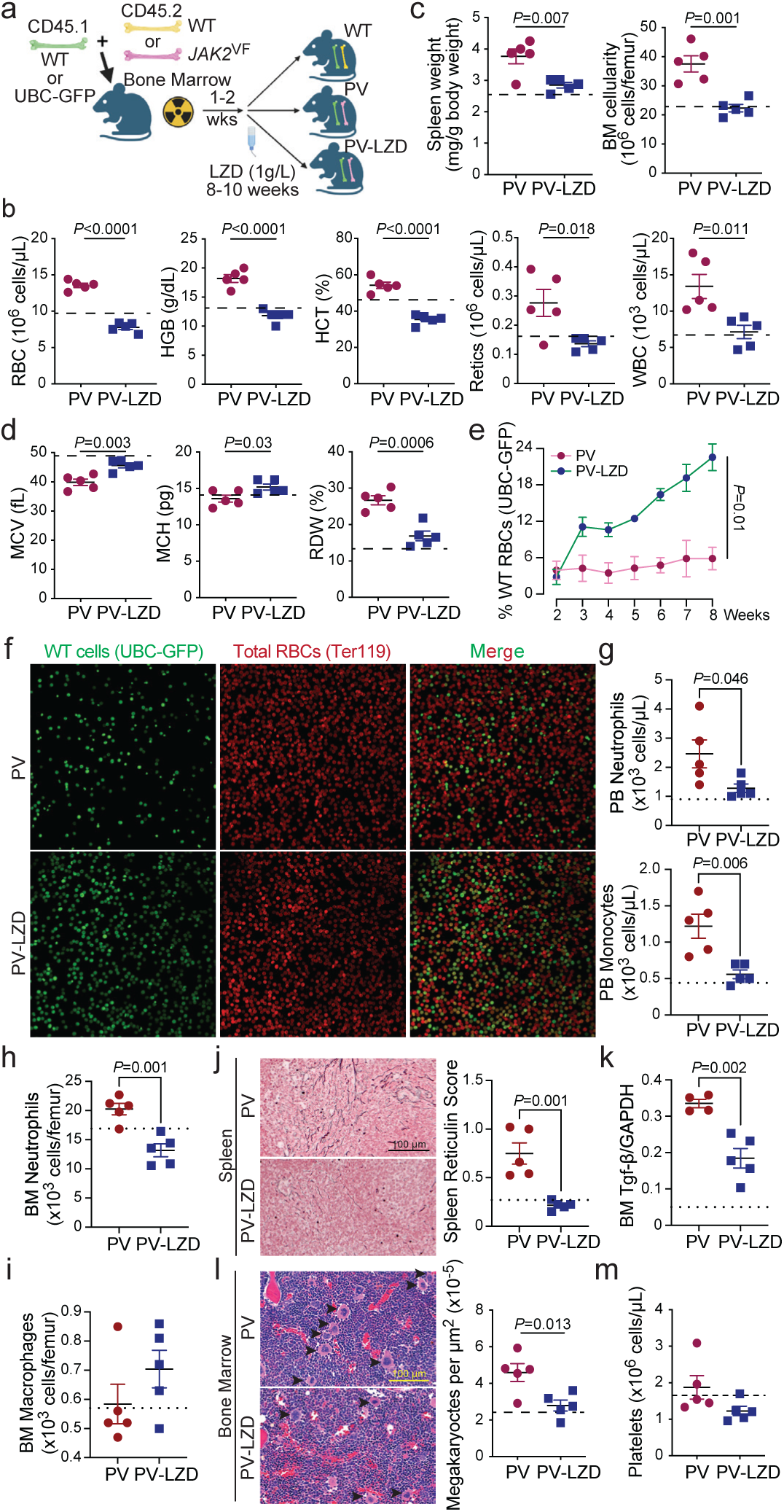
LZD selectively inhibited *JAK2*^VF^ cells and reduced splenic fibrosis. Mice were irradiated with X-ray at a dose of 1.2 Gy before bone marrow transplantation. C57BL/6 mice received CD45.1 and CD45.2 grafts from wild-type (WT) mice, whereas polycythaemia vera (PV) mice received CD45.1 grafts from WT (or UBC-GFP, panels e and f) mice and CD45.2 grafts from *JAK2*^VF^ mice. Irradiated PV mice were divided into two groups, half of which were treated with LZD (1 g/L in drinking water) for 8–10 weeks (**a**). Shown are peripheral blood counts, namely red blood cell (RBC), haemoglobin (HGB), haematocrit (HCT), reticulocytes (Retics) and white blood cell (WBC) (**b**), normalized spleen weight and bone marrow cellularity (**c**), mean corpuscular volume (MCV), mean corpuscular haemoglobin (MCH) and red cell distribution width (RDW) (**d**), percentage of WT (UBC-GFP) cells (**e**), distribution of WT (UBC-GFP) and RBC (Ter119+) cells (**f**), and neutrophil and monocyte counts (**g**) in peripheral blood. Also shown are neutrophil (**h**) and macrophage (**i**) counts in bone marrow (**j**), splenic fibrosis (reticulin stain) (**j**) expression (normalzied to GAPDH) (**k**), bone marrow megakaryocytes (arrows) (**l**), and platelet counts (**m**) in PV mice treated with (PV-LZD) or without LZD. For each panel, the dash line indicates mean levels in the WT group. Scale bar = 100 µm (j). Statistics: mean ± s.e.m., *N*=5 mice per group, unpaired two-tailed Student’s *t*-test, *P* values shown.

### LZD Selectively Inhibits *JAK2*^VF^ Cells, and Lowers the *JAK2*^VF^ Allele Frequency, Proinflammatory Cells, Megakaryocyte Counts and Fibrosis in Competitively Transplanted Mice

Given that the bone marrow of PV patients contains healthy normal hematopoietic cells along with mutated *JAK2*^VF^-expressing cells^25,26^, we utilized a competitive bone marrow transplantation strategy where bone marrow cells from WT (CD45.1) and *JAK2*^VF^ (CD45.2) mice were mixed and transplanted into lethally-irradiated recipient mice. The WT group received mixed WT (CD45.1) and WT (CD45.2) grafts; the PV group received WT (CD45.1) plus *JAK2*^VF^(CD45.2) grafts; and the PV-LZD group received WT plus *JAK2*^VF^ grafts and was treated with 1 g/L LZD in drinking water (Figure 5A). Mice from the PV group expectedly developed erythrocytosis, leukocytosis, splenomegaly and increased marrow cellularity (Fig 4B, C). Remarkably, LZD treatment for 8 weeks led to reduced hematocrit, leukocyte counts, and splenomegaly, and normalized marrow cellularity (Figure 5B, C). Of note is that our competitive PV model contained both WT and *JAK2*^VF^ mutant cells. LZD treatment significantly suppressed white blood counts from both WT and *JAK2*^VF^ populations compared to PV group (supplemental Figure 2). LZD increased the RBC mean corpuscular volume and mean corpuscular hemoglobin, and decreased red cell distribution width, shifting these parameters towards levels seen in WT mice (Figure 5D).

As CD45.1 and CD45.2 markers are not expressed on erythroid cells, and to confirm that LZD promoted the release of WT red blood cells, while restricting *JAK2*^VF^ red blood cell release, we performed competitive bone marrow transplants with a mix of WT UBC-GFP cells and unmarked *JAK2*^VF^ mutant cells. Since the number of circulating red blood cells is about 1000x greater than white blood cells in peripheral blood, UBC-GFP fluorescence signals arose mainly from red blood cells. After 8 weeks of LZD treatment, mice progressively increased the percentage of circulating WT UBC-GFP cells, while *JAK2*^VF^ cells were reduced (Figure 5E). Peripheral blood smears further showed more WT UBC-GFP+ cells in the PV-LZD group compared to the PV group, with a comparable density of total red blood cells (Ter119+) between the two groups (Figure 5F). These data demonstrated, for the first time, that LZD specifically suppressed the dominant *JAK2*^VF^ red blood cells in the peripheral blood and promotes WT red blood cells release—as evidence for a profound reduction of *JAK2*^VF^ allele frequency in WT plus *JAK2*^VF^ grafted mice.

Increased numbers of proinflammatory cells, namely neutrophils and monocytes/macrophages, in PV patients promote inflammation and are predictive of poor outcomes^27–29^. We found that LZD reduced peripheral blood neutrophils and monocytes (Figure 5G), and neutrophil numbers in the bone marrow (Figure 5H). There were no differences in bone marrow macrophage numbers (Figure 5I). LZD also reduced the degree of in fibrosis in the spleens of the PV group (Figure 5J). No bone marrow fibrosis was observed in the bone marrow of either PV or PV-LZD groups (supplemental Figure 3). Bone marrow megakaryocyte numbers were reduced in the PV-LZD group (Figure 5K). Quantitative PCR nonetheless showed that LZD reduced the expression of *Tgfb*, a known mediator of myelofibrosis (Figure 5L). The upregulated megakaryocytes count did not translate into elevated platelet production in the PV group (Figure 5M). Thus, platelet counts remained unchanged in both the PV and PV-LZD groups (Figure 5M)^28–32^.

### LZD Selectively Reduces Disease-Initiating *JAK2*^VF^ Stem Cells in Transplanted Mice

*JAK2*^VF^ mutated stem cells, especially long-term hematopoietic stem cells (LT-HSC), are known to be the disease-initiating cells in PV^4^. We used gating strategies to assess the impact of LZD on hematopoietic stem cell and progenitor compartments in our competitive bone marrow transplant model (supplemental Figure 4A). LZD treatment normalized the total number of LT-HSCs compared to the PV group (supplemental Figure 4B). Importantly, the PV-LZD group displayed a reduction in the expanded WT LT-HSC numbers (CD45.1) *vs.* the PV group. Most notable was a clear reduction in the number of disease-inducing LT-HSC *JAK2*^VF^ cells (CD45.2) (Figure 6A). Of note is t the observed expansion of the WT fraction within the LT-HSC compartment in the PV group which was consistent with previous studies showing that co-culture of WT and *JAK2*^VF^ hematopoietic stem and progenitor cells promoted WT expansion *vs*. WT cells cultured alone^33^. In sum, these findings highlight the ability of LZD to directly target and deplete the mutated HSC compartment.

**Figure 6:**
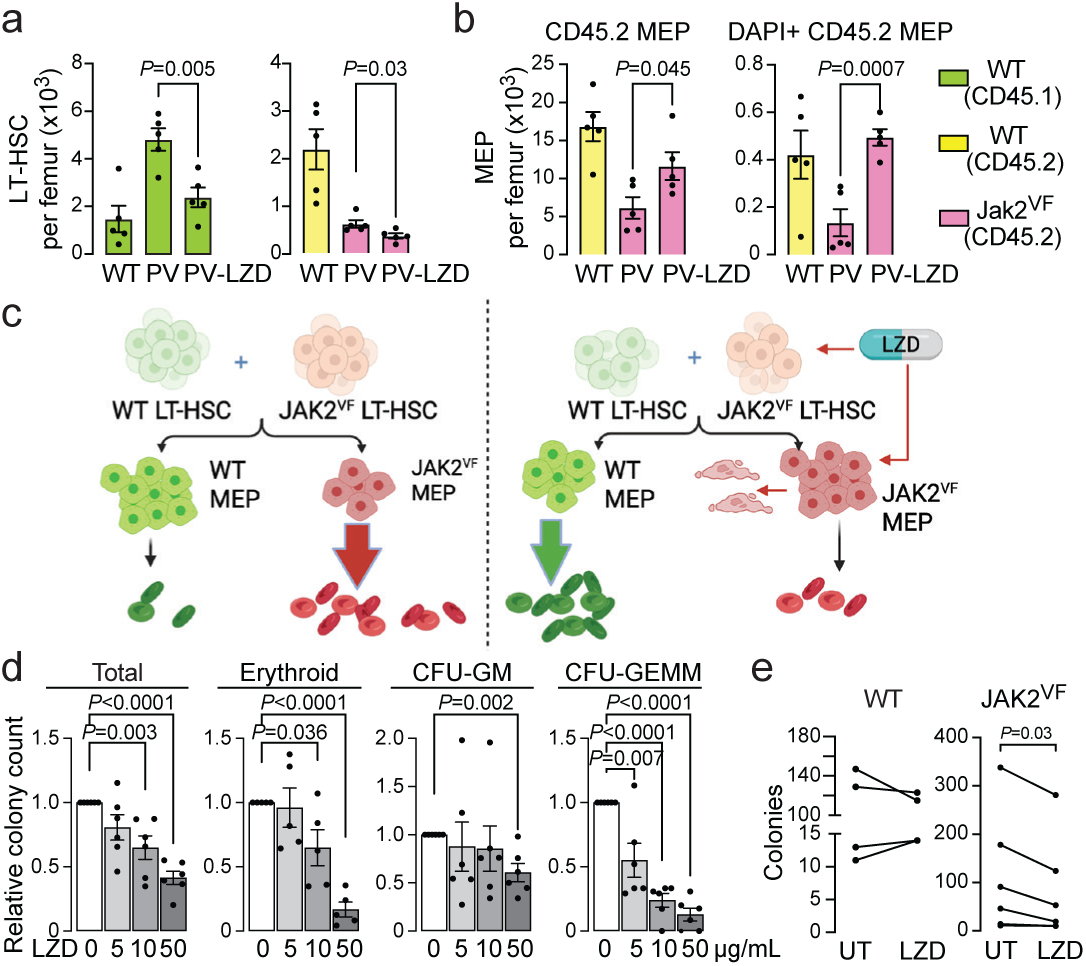
LZD treatment depleted JAK2^VF^ stem cell in mice and suppressed haematopoietic progenitors in PV patients. Irradiated C57BL/6 mice received wild type (WT) CD45.1 and CD45.2 grafts, whereas polycythaemia vera (PV) mice received WT CD45.1 and *JAK2*^VF^ CD45.2 grafts. A group of PV mice received LZD (1 g/L in drinking water) for 8–10 weeks. Shown are bone marrow long-term haematopoietic cells (LT-HSC) (**a**) and total CD45.2 megakaryocyte-erythroid progenitor (MEP) and dead DAPI+ CD45.2 MEP cells (**b**). (**c**) Working model: Compared to untreated (Ctrl) group, LZD reduced LT-HSC number and promoted cell death in erythroid lineages which further decreased release of JAK2^VF^ cells and increased WT cells in circulation. (**d**) Total colony count, erythroid colonies (CFU-E + BFU-E), Granulocyte-macrophage (CFU-GM) and granulocyte-erythrocyte-monocyte-mega-karyocyte (CFU-GEMM) in PV patient treated with LZD (5, 10 or 50 µg/mL). (**e**) Number of WT- and JAK2^VF^-containing colonies from PV patients treated or untreated (UT) with LZD. Statistics: mean ± s.e.m., *N*=5 mice (a, b) and 6 patients (d, e) per group, unpaired two-tailed Student’s *t*-test, *P* values shown.

The megakaryocyte-erythroid progenitor population was also altered with LZD treatment. While the total number of megakaryocyte-erythroid progenitor was similar between PV and PV-LZD groups (supplemental Figure 4C), there were lower levels of *JAK2*^VF^ megakaryocyte-erythroid progenitor cells in PV group, whereas the *JAK2*^VF^ megakaryocyte-erythroid progenitor cells in the PV-LZD group were increased (Figure 6B). WT-derived megakaryocyte-erythroid progenitor cells showed no difference between the PV and PV-LZD groups (supplemental Figure 4D). As summarized in our working model, despite the accumulation of *JAK2*^VF^ megakaryocyte-erythroid progenitor cells in PV-LZD group, LZD promoted cell death in this population indicated by upregulating DAPI⁺ cells (Figure 6C, supplemental Figure 4E). We also found fewer *JAK*2^VF^ megakaryocyte-erythroid progenitor cells in bone marrow of the PV group, which was associated with greater release of *JAK*2^VF^ erythroid cells in peripheral blood (Figure 4E, F, 6C). In contrast, LZD altered the dynamics of cell retention in the bone marrow, such that more *JAK2*^VF^ mutant megakaryocyte-erythroid progenitor cells accumulated and underwent cell death, with fewer *JAK2*^VF^ erythroid cells being released into peripheral blood (Figure 4E, F, 6C).

### LZD Suppresses Hematopoietic Colony Formation by PV Patient Mononuclear Cells

To evaluate if LZD affects human PV progenitor cells, we performed hematopoietic colony assays with peripheral blood mononuclear cells from six PV patients. LZD reduced colony formation in a concentration-dependent manner (Figure 6D). The colonies included erythroid colonies (BFU-E and CFU-E), colony-forming unit-granulocyte/macrophage (CFU-GM), and colony-forming unit-granulocyte/erythrocyte/macrophage/megakaryocyte (CFU-GEMM), with CFU-GEMM representing the most primitive progenitor population. Erythroid colony formation was suppressed at doses of 10 and 50 µg/mL LZD, whereas CFU-GM colony number was reduced at 50 µg/mL (Figure 6D). CFU-GEMM colonies were reduced significantly with LZD at a concentration as low as 5 µg/mL with further reductions with increasing concentrations (Figure 6D). These data collectively indicated that LZD preferentially targets primitive human progenitor cells forming CFU-GEMM colonies at the low concentrations. Most importantly, genotyping of individual colonies revealed that LZD selectively reduces the number of JAK2^VF+^ colonies (Figure 6E), while there was no impact on the reservoir of WT colonies (Figure 6E)—yet again demonstrating the specificity of LZD in targeting JAK2^VF+^ cells in PV patient samples.

## DISCUSSION

The pursuit of JAK2^VF^IZselective inhibitors has gained importance in recent years, with multiple groups attempting to develop agents that preferentially suppress mutant signaling while sparing wild type JAK2^34,35^. These preclinical datasets show that these selective inhibitors exhibit markedly greater potency against JAK2^VF^IZmutant cells than WT cells in both murine models and human cellIZbased assays. Two groups^34,35^ recently presented earlyIZstage data in abstract form on reducing JAK2^VF^ cells, further underscoring the growing interest in this area. Although these inhibitors demonstrate preferential activity against JAK2^VF^IZmutant cells, no study to our knowledge has yet established that WT hematopoiesis remains intact or, indeed, that PV hematopoietic stem cells are inhibited. By contrast, our findings clearly document that LZD suppresses proliferation across multiple *JAK2*^VF^IZdriven systems—including *JAK2*^VF^ mice, HEL cells, and PV patient progenitor cells—and acts primarily through a direct interaction with stem/progenitor cell populations to ultimately reduce variant allele frequency. Even more important is that LZD produces no detectable hematopoietic toxicity in WT mice when administered at doses sufficient to rescue *JAK2*^VF^IZassociated phenotypes, supporting a favourable safety profile. The convergence of efficacy across these models suggests a targeted mechanism of action, reinforcing the therapeutic potential of LZD and future LZDIZlike compounds as mutantIZdirected interventions for JAK2^VF^IZdriven hematologic disorders.

Reinforcing the selectivity of LZD, we provide a clear mechanism for LZD action on the JAK2^VF^ mutation, while sparing WT JAK2. The disease-causing JAK2^VF^ mutation results from a V617F substitution in the pseudokinase, JH2, domain of the JAK2 protein^36,37^. In the JAK2^VF^ protein, residues F617 at the mutation site and F595 in the unmutated region form a potential π stacking interaction that stabilizes an active conformation and promotes autophosphorylation of the JAK2^VF^ protein^36,37^. Disrupting this F617F–F595 interaction abolishes JAK2^VF^ autophosphorylation, underscoring its functional importance^36,37^. Current JAK2 inhibitors, such as ruxolitinib, target the JAK2 tyrosine kinase (JH1) domain, not the pseudokinase (JH2) domain, leading to the widespread and indiscriminate inhibition of mutated as well as JAK2 proteins^24,38^. Our data instead show that LZD engages the F617–F595 interface by docking in between these residues to suppress JAK2/STAT5 signalling––thus offering a novel, highly specific, first-in-class therapeutic for selectively blocking JAK2^VF^ activity through direct targeting of the JH2 domain.

## ACKNOWLEDGEMENTS

We express our deepest gratitude to the late Dr. Paul Frenette for his invaluable guidance. This work was supported by the National Institute of Diabetes and Digestive and Kidney Diseases (NIDDK) (K01DK131401 to H.L.); the National Cancer Institute (P01CA108671 to R.H.); the National Institute on Aging (R01AG071870, R01AG074092, and U01AG073148 to T.Y. and M.Z.; and U19AG060917 to M.Z.); the National Heart Lung and Blood Institute (R01HL159116 and U01HL167036 for REAL Answers to J.G); the National Institute of Diabetes and Digestive and Kidney Diseases (R01DK143032, R01DK107670, R01DK095112, to Y.Z.; R56DK132146 to Y.Z. and R.H; and K01DK137045 to X.G.); the MPN Foundation (Y.Z.); the ASH Fellow-to-Faculty Scholar Award, and V Scholar Grant from V Foundation for Cancer Research to X.G. the Hevolution Foundation supported M.Z. and T.Y.

## AUTHORSHIP CONTRIBUTIONS

A.A.G. and C.S. performed the experiments and wrote the manuscript. Z.H., A.A., S.H., A.P., J.K., F.F.G., J.C., G.P., C.Y., Z.T. and S.L. participated in experiments. X.G. participated in experiments and revised the manuscript. T.Y., P.F., J.G., S.H., Y.G. revised the manuscript. M.Z. designed the molecular docking experiment and revised the manuscript. R.H. designed the human patient sample experiments, analysed the data and revised manuscript. H.L. designed and performed all experiments and wrote the manuscript.

## DISCLOSURE OF CONFLICTS OF INTEREST

Authors Huihui Li, Ronald Hoffman, Mone Zaidi, and Shozeb Haider are inventors on a patent application related to repurposing LZD for treatment of diseases with Jak2^VF^ mutation, which has been filed by the Icahn School of Medicine at Mount Sinai. All other authors declare no competing interests.

---

**supplemental Figure 1.**
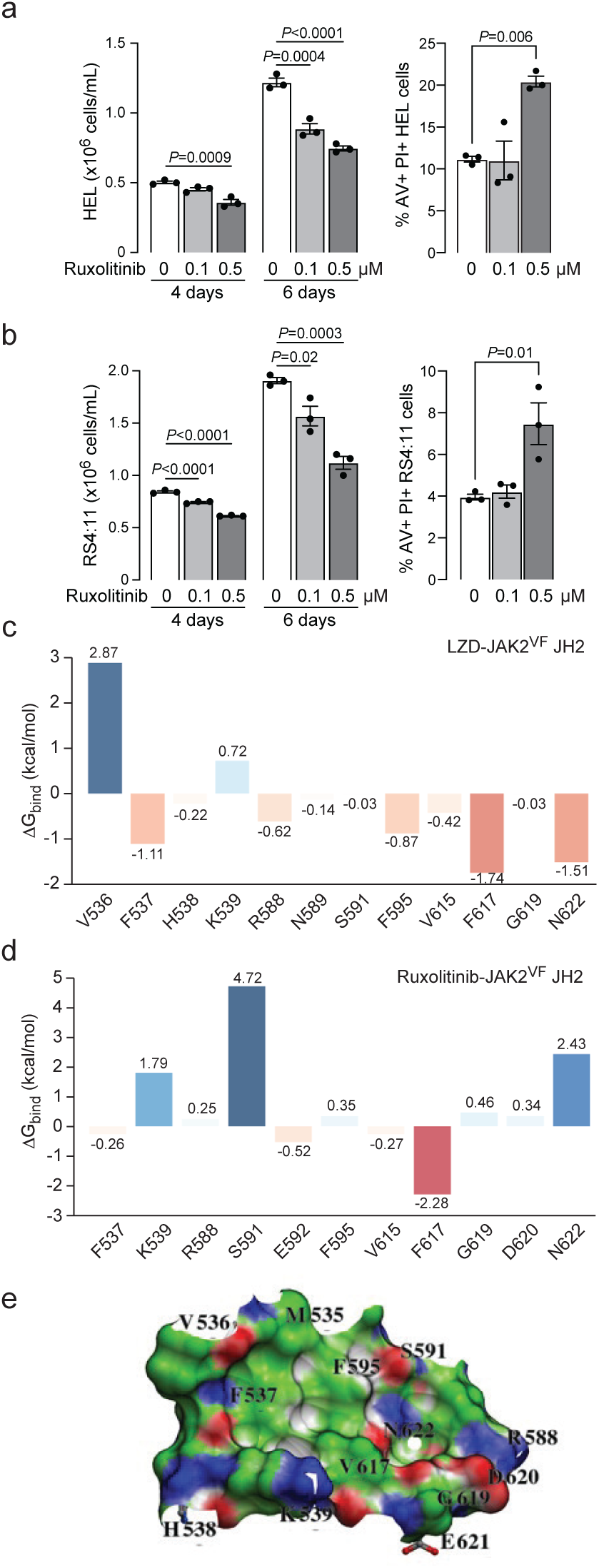
: Unlike LZD, ruxolitinib inhibited both WT and mutant JAK2^VF^. Treatment of ruxolitinib suppressed cell proliferation and promoted apoptosis of HEL (**a**) and RS4:11 (**b**) cells. Per-residue contribution to binding energy of LZD (**c**) and Ruxolitinib (**d**) in JH2 domains. LZD does not bind to the JH2 domain of WT JAK2 protein (**e**). Statistics (a, b): mean ± s.e.m., *N*=3 per group, unpaired two-tailed Student’s *t*-test (vs. 0 µg/mL LZD), *P* values shown.

**supplemental Figure 2.**
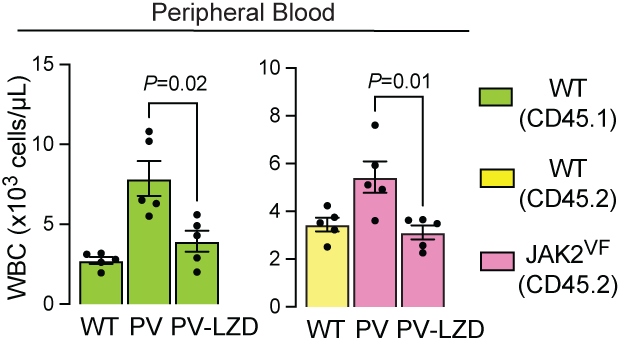
: Linezolid (LZD) reduced WBC release from bone marrow. Irradiated C57BL/6 mice received wild type (WT) CD45.1 and CD45.2 grafts, whereas polycythaemia vera (PV) mice received WT CD45.1 and *JAK2*^VF^ CD45.2 grafts. A group of PV mice received LZD (1 g/L in drinking water) for 8–10 weeks. Shown are peripheral WBC counts following treatment.

**supplemental Figure 3.**
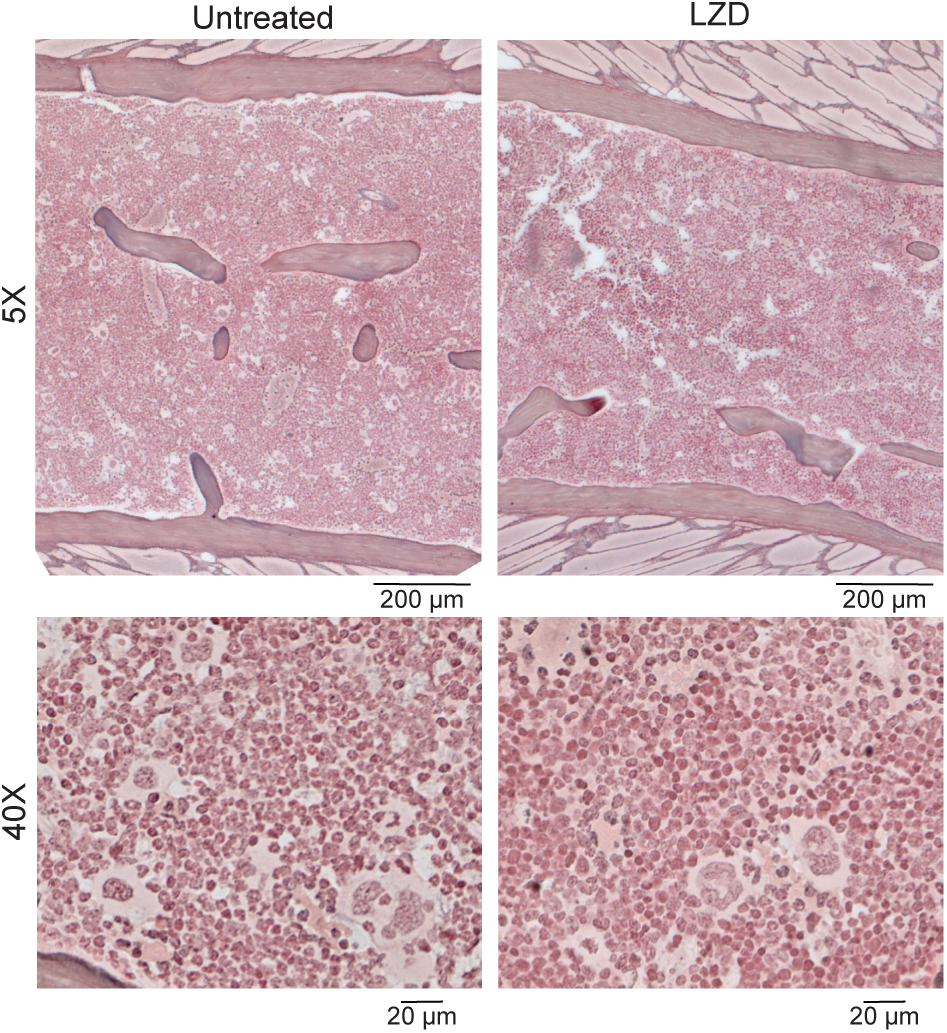
: Marrow fibrosis (Reticulin staining) was not detected in competitive bone marrow transplantated (WT+JAK2^VF^) mice in both untreated and LZD groups. Scale bar = 200 µm (top panels), 20 µm (bottom panels).

**supplemental Figure 4 :**
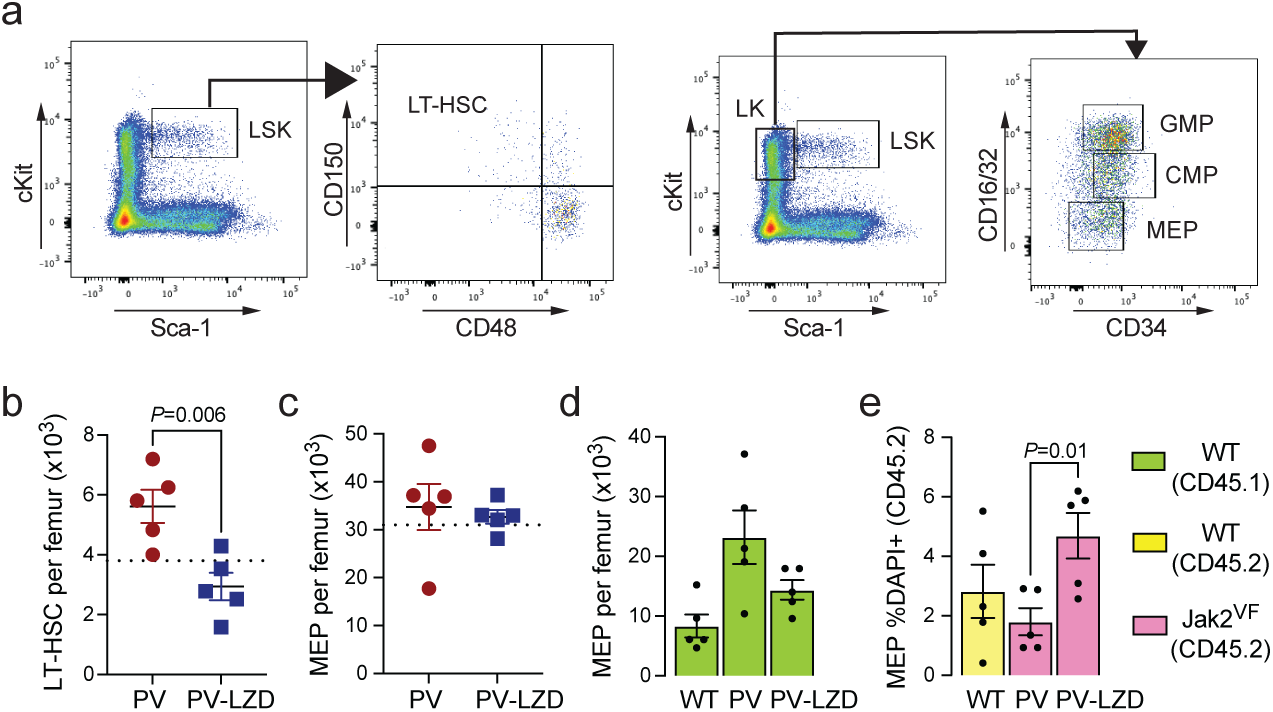
LZD impacted JAK2^VF^ stem cell numbers in competitive BMT (WT+*JAK2*^VF^) mice. (**a**) Gating strategies for haematopoietic stem cells; Total number of (**b**) long-term haematopoietic stem cells (LT-HSC), and (**c**) megakaryocyte-erythroid progenitor (MEP) from PV and PV-LZD groups; Dashed line represented levels in WT group; (**d**) Absolute number of CD45.1 MEP cells from different groups; and (**e**) percentage of CD45.2 MEP cells from different groups. Statistics (b-e): mean ± s.e.m., *N*=5 mice per group, unpaired two-tailed Student’s *t*-test, *P* values shown.

## Notes

### Competing Interest Statement

The authors have declared no competing interest.

