## Supplemental method for "Linezolid Acts as a Selective Inhibitor of the JAK2^V617F^ Mutation"

**Bone marrow transplantation**

Recipient C57BL/6 mice at 8 weeks were subjected to total body irradiation (TBI) at a dose of 1.2 Gy, delivered as a split dose (0.6 Gy twice, 11 hours apart) using an X-ray irradiator (Precision). For animals exclusively receiving bone marrow transplants, 2 million JAK2^VF^ bone marrow cells were administered via retro-orbital injection. For competitive bone marrow transplants with CD45.1 and CD45.2 cells, irradiated mice receive 1 million CD45.1 (WT) and 1 million CD45.2 (JAK2^VF^ or WT) bone marrow cells. For competitive bone marrow transplantation with UBC-GFP and JAK2^VF^ cells, irradiated mice receive 1 million UBC-GFP and 3 million JAK2^VF^ bone marrow cells. Recipient mice were housed in clean cages and provided with autoclaved food and acidified water. Blood samples from the tail vein were taken at time intervals ranging from 1 to 12 weeks after transplantation to monitor the degree of chimerism.

**Flow cytometry**

Bone marrow and peripheral blood cells were harvested and washed once with phosphate-buffered saline (PBS). Following the lysis of red blood cells using ammonium chloride-potassium (ACK) buffer, marrow cells were resuspended in flow cytometry buffer (PBS supplemented with 0.5 % BSA and 2 mM EDTA) on ice. For phenotypic analyses, 1 x 10^6^ cells were incubated with fluorophore-conjugated monoclonal antibodies (as listed in Supplemental Table 1) for 0.5–1 hour at 4°C in the dark. Cells were then washed twice with flow cytometry buffer and stained with 4’,6-diamino-2-phenylindole (DAPI) and analysed immediately. Neutrophils were identified as Gr-1^hi^CD115^lo^SSC^hi^ cells. T cells, B cells, and monocytes were gated as CD3⁺CD11b^–^, B220⁺CD11b^–^, and CD115^hi^CD11b⁺ populations, respectively. Haematopoietic stem and progenitor cells (HSPCs) were defined as lineage-negative (Gr-1^–^, CD11b^–^, CD3^–^, B220^–^, and Ter119^–^) and further characterized by Sca-1, c-Kit, CD135, CD150, CD48, CD34, and CD16/32 expression (supplemental Figure 4a, supplemental Table 2). Macrophages were identified as Gr-1^lo^CD115^lo^F4/80⁺SSC^lo^ cells. Erythroblasts were identified as CD44^hi^/FSC^hi^ cells. Enucleated cells were defined by CD44^med/low^/FSC^low^. Cell cycle was analysed using Ki67 and DAPI. Data were acquired on a Symphony A5 flow cytometer (BD Biosciences) and analysed using FlowJo software (v.10.4).

*In vitro* experiment, apoptotic cells were identified with annexin-V/propidium iodide staining. Cells were harvested, washed twice with cold PBS, and resuspended in 1x annexin V binding buffer (10 mM HEPES, 140 mM NaCl, 2.5 mM CaCl_2_, pH 7.4). Annexin V conjugated to fluorescein isothiocyanate (annexin V–FITC) and propidium iodide (PI) were added *per* manufacturer’s instructions (ThermoFisher Scientific) (supplemental Table 2). Samples were incubated for 15 minutes at room temperature in the dark. Immediately following incubation, an additional binding buffer was added, and data were acquired on a Symphony A5 flow cytometer and analysed using FlowJo (v.10.4). Live cells were identified as annexin V⁻/PI⁻, early apoptotic cells as annexin V⁺/PI⁻, and late apoptotic/necrotic cells as annexin V⁺/PI⁺. *In vivo* experiment, dead cells from spleen and bone marrow samples were identified as DAPI^+^ cells. FACS analyses were carried out using LSR II flow cytometry (BD Biosciences). Data were analysed with FlowJo v.10.4 and FACS Diva v.6.1.

**Hematological Analyses**

Blood was harvested from the retro-orbital plexus into heparinized tubes and preserved in 6 µL 0.5 M EDTA. Complete blood counts, white blood cell differentials, and reticulocyte counts were performed using an ADVIA 120 Hematology System (Siemens).

**Histology**

*Tissue collection and processing*: Spleens and sternums were harvested, fixed in 10% neutral-buffered formalin (NBF) for 24 hours at room temperature, and rinsed in PBS. Sternums were decalcified using Morse’s solution at room temperature for 6–8 hours. Tissues were then processed using a standard paraffin embedding protocol. Paraffin sections were made with a thickness of 4 µm.

*H&E staining*: Standard protocols were used. Prussian blue stain was performed with Iron Stain Kit (StatLab, cat.# KTIRO). Reticulin fibres were visualized by silver impregnation using a Chandler’s Precision Reticulum Stain Kit (StatLab, cat.# KTCPR) according to manufacturer’s protocols.

*Microscopic Analysis*: Slides were examined under a brightfield microscope (Zeiss) at 10x and 20x magnifications. The spleen was evaluated for the presence of white pulp (lymphoid follicles and germinal centres) and red pulp (sinusoids and splenic cords). Megakaryocytes were identified based on their characteristically enlarged morphology, and their numbers were normalized to the bone‑marrow area.

**RNA Isolation and Quantitative Real-Time PCR of bone marrow cells**

Total RNA was extracted from bone marrow cells using the RNeasy Mini Kit (Qiagen) according to the manufacturer’s protocol. cDNA synthesis was performed using the High-Capacity cDNA Reverse Transcription Kit (Applied Biosystems) with 2 µg of total RNA per reaction. Quantitative real-time PCR (RT-qPCR) was carried out using PowerUp SYBR Green Master Mix (ThermoFisher Scientific) on a QuantStudio 6 Flex Real-Time PCR System (Applied Biosystems). Gene-specific primers targeting inflammatory cytokine *Tgfb* was designed using Primer-BLAST and synthesized by Integrated DNA Technologies (IDT). *Tgfb* primer: Forward: TGATACGCCTGAGTGGCTGTCT, Reverse: CACAAGAGCAGTGAGCGCTGAA. Relative expression levels were calculated using the ΔΔCt method with *Gapdh* as the endogenous control. All reactions were performed in technical triplicates and melt curve analysis was conducted to confirm amplification specificity.

**Randomization and Statistics**

Mice were randomly assigned to experimental groups using a simple randomization procedure to ensure an unbiased distribution of baseline characteristics, including body weight. Each data point is from a distinct sample. Statistically significant differences between any two groups were examined using a unpaired two-tailed Student’s *t*-test, given equal variance. P values were considered significant at or below 0.05.

**Data collection and availability**

Microsoft Excel and GraphPad Prism (v.10) are used to collect data for this study. The authors declare that all data supporting the findings of this study are available within the paper and as Source Data files. Specifically, all Source Data for Figs 1, 2, 3, 4, 5, 6 and Supplemental Figs 1, 2, 3, 4 is available with the online version of the paper as a PDF file. Supplemental Table 1 attributes the antibody lists for experiments.
