## Supplemental table for "Linezolid Acts as a Selective Inhibitor of the JAK2^V617F^ Mutation"

**supplemental Table 1:** Patient information

| **Patient** | **Sex** | **Age at sampling (yrs)** | **Disease** | **Driver mutation (VAF)** | **Other myeloid-associated mutations (VAF)** | **Karyotype/aCGH+SNP** |
| --- | --- | --- | --- | --- | --- | --- |
| 1 | M | 55 | PV | JAK2-V617F (32.3%) | none | 46,XY[20] |
| 2 | M | 59 | PV | JAK2-V617F (39.0%) | IDH2-R140Q (5%) | 46,XY[20]/ arr[GRCh37] 3p22.3p22.1(34868770_39871369)x2 hmz, arr[GRCh37] 8q24.11q24.13(119085415_124938907)x2 hmz, arr[GRCh37] 9p24.3p21.2(0_27550437)x2 hmz |
| 3 | F | 60 | pPV-MF | JAK2-V617F (89.1%) | NRAS-G12D (10.0% SH2B3-K181* (20.6%) | 46,XX,dup(1)(q21q42)[1]/46,XX[19] |
| 4 | M | 66 | PV | JAK2-V617F (61.6%) | not available | not performed |
| 5 | M | 35 | PV | JAK2-V617F (77.3%) | ASXL1-G646Wfs*12 (30.9%) EZH2-E173Q (29.3%) | 46,XY[20] |
| 6 | F | 57 | PV | JAK2-V617F (84%) | KRAS-G12R (7%) TET2-Y1245Lfs*22 (39%) TET2-Q876* (46%) | not available |

**supplemental Table 2:** List of antibodies used.

| **Reagent** | **Catalog No.** | **Manufacturer** |
| --- | --- | --- |
| Anti-mouse TER119 | 116208 | BioLegend |
| Anti-mouse CD44 | 20-0441-U100 | TonBo Biosciences |
| Anti-mouse CD71 | 113820 | BioLegend |
| Anti-mouse CD71 | 113806 | BioLegend |
| Anti-mouse CD71 | 25-0711-82 | eBioscience |
| Anti-human CD45 | 304038 | BioLegend |
| Anti-human/mouse p-STAT 3 | 11-9033-42 | Invitrogen |
| Anti-human/mouse p-STAT 5 | 12-9010-42 | Invitrogen |
| Anti-mouse p-ERK | 675504 | BioLegend |
| Anti-mouse CD45.1 | 110737 | BioLegend |
| Anti-mouse CD45.1 | 47-0453-82 | BioLegend |
| Anti-mouse CD45.1 | 110714 | BioLegend |
| Anti-mouse CD45.2 | 109835 | BioLegend |
| Anti-mouse CD45.2 | 109824 | BioLegend |
| Anti-mouse CD11b | 25-0112-82 | Invitrogen |
| Anti-mouse CD11b | 45-0112-82 | Invitrogen |
| Anti-mouse Ki-67 | 652403 | BioLegend |
| Anti-mouse CD117 (c-Kit) | 105828 | BioLegend |
| Anti-mouse Ly-6A/E (Sca-1) | 11-5981-82 | eBioscience |
| Anti-mouse Ly-6A/E (Sca-1) | 25-5981-82 | eBioscience |
| Anti-mouse Ly-6A/E (Sca-1) | 17-5981-83 | eBioscience |
| Anti-mouse F4/80 | 123107 | BioLegend |
| Biotin Mouse Lineage Panel | 559971 | BD Biosciences |
| Anti-mouse CD48 | 103416 | BioLegend |
| Anti-mouse CD3 | 100203 | BioLegend |
| Anti-mouse Gr-1 | 108405 | BioLegend |
| Anti-mouse CD115 | 165003 | BioLegend |
